# The muscle atrophic phenotype of MuSK myasthenia gravis: Insights from a preclinical rat model

**DOI:** 10.64898/2026.03.13.709755

**Authors:** Jesper Emil Jakobsgaard, Pernille Bogetofte Thomasen, Jakob Wang, Thea Holm Kristiansen, Pernille Johnsen, Anders Riisager, Nete Huus, Martin Broch-Lips, Martin Brandhøj Skov, Johan Palmfeldt, Thomas Holm Pedersen, Kristian Vissing

## Abstract

Myasthenia gravis with muscle-specific kinase antibodies (MuSK-MG) is an autoimmune disorder marked by neuromuscular junction (NMJ) disruption and selective muscle atrophy, yet its intrinsic myocellular mechanisms remain unclear. Using a rat model generated by active immunization with the N-terminal MuSK60 peptide, we characterized muscle pathology, NMJ morphology, and whole-muscle proteome remodeling across anatomically distinct skeletal muscles. Anti-MuSK rats developed seropositivity, body-mass loss, fragmented and denervated NMJs, and pronounced atrophy restricted to slow-twitch/type I fibers, particularly in the soleus muscle. Quantitative proteomics identified extensive muscle-specific alterations, most prominent in soleus, with fewer in diaphragm and sternohyoideus. Gene set enrichment analysis revealed coordinated downregulation of mitochondrial, ribosomal, and myosin-complex proteins in soleus, partial mitochondrial involvement in diaphragm, and compensatory upregulation of translational and proteasomal pathways in diaphragm. Correlation analysis linked soleus mass loss to elevated abundance of ubiquitin-proteasome and calcium-handling proteins, implicating proteolytic and bioenergetic stress mechanisms. The consistent upregulation of NCAM1 and MUSTN1 suggests generalized myocellular responses to MuSK dysfunction. Together, these data demonstrate that MuSK autoimmunity elicits fiber-type–selective atrophy and profound proteome remodeling beyond NMJ impairment, highlighting disrupted mitochondrial and translational homeostasis as central features of the MuSK-MG muscle phenotype.

## 1. Introduction

Myasthenia gravis (MG) is a rare autoimmune neuromuscular disorder characterized by fluctuating muscle weakness and fatigue (1–3). While the pathogenesis of MG is primarily attributed to autoantibodies targeting the acetylcholine receptor (AChR) at the neuromuscular junction (NMJ), a subset of MG patients does not exhibit these antibodies (1,4–8). Instead, a majority of this subset exhibit circulating antibodies against the muscle-specific kinase (MuSK), an MG subtype discovered as recently as in the early 2000s (7–9). Given its well-described role in developing and stabilizing AChR-clusters at the neuromuscular junction, impaired MuSK function mediated by the antibody attack, results in weakened neuromuscular transmission similar to AChR-MG (10–12). However, the pathophysiology and therapeutic responses of the MuSK subtype differ from AChR-MG (13–16), underscoring the need for tailored approaches in diagnosis and treatment.

In MuSK-MG patients, muscle atrophy is reported as a distinct clinical observation in various affected muscles, mainly in tongue as well as other facial and bulbar muscle groups (5,8,17–21), extrinsic ocular (22), and paraspinal and limb muscles (23,24). Moreover, muscle atrophy is a feature in experimental animal models of MuSK-MG (25–28). The underlying mechanisms of this muscle-atrophic disease trait may entail multiple signaling components. Developmental maturation and postnatal stability of the NMJ is largely dependent on the canonical signaling cascade by which MuSK is activated through the neural secreted proteoglycan Agrin and its receptor, the low-density lipoprotein receptor-related protein 4 (LRP4) (10–12). Additionally, additional signaling roles for MuSK have emerged within the last decades, pointing toward its involvement in myocellular regulatory events beyond canonical signaling in NMJ formation and integrity. For example, MuSK has recently been implicated as a co-receptor in transforming growth factor beta (TGF-β) signaling, a superfamily important for regulating muscle mass (29–31). Thus, disruption of MuSK function may introduce an active muscle-atrophic component besides impaired neuromuscular transmission in the affected muscles of MuSK-MG patients.

Unraveling the molecular properties of the muscle-atrophic disease trait in MuSK-MG has important implications for the understanding of disease etiology. The impact of the MuSK-MG pathogenesis on neuromuscular transmission parameters and synaptic properties at the NMJ has been studied and described in both patient cohorts and animal models (3,14–16,18,21,25,27,32–36). However, the intrinsic myocellular mechanisms and properties of MuSK-MG are not well-described leaving important knowledge gaps concerning the myocellular phenotype of MuSK. Consequently, using whole-muscle proteomics and immunohistochemical approaches, the aim of the present study was to conduct in-depth characterizing of the myocellular phenotype in a preclinical model of the MuSK-MG pathogenesis.

## 2. Methods

### Ethical approval

All experimental procedures were approved by the Danish Animal Experimental Inspectorate (license no. 2018-15-0201-01420). The study was conducted in concordance with Danish Animal welfare legislation and the Directive 2010/63/EU of the European Parliament and the Council of 22 September 2010.

### Study design

To establish a cross-sectional design for studying the MuSK-MG phenotype (Figure 1A), 18 rats were enrolled in the present study and allocated to either an Anti-MuSK group receiving a MuSK antigen (see Disease model section, n=12) or Sham control group (n=6). Additionally, 1 untreated rat was euthanized as a muscle donor for staining of wild-type muscle sections (see “Serum MuSK antibody staining in wild-type rat muscle” section).

**Figure 1.**
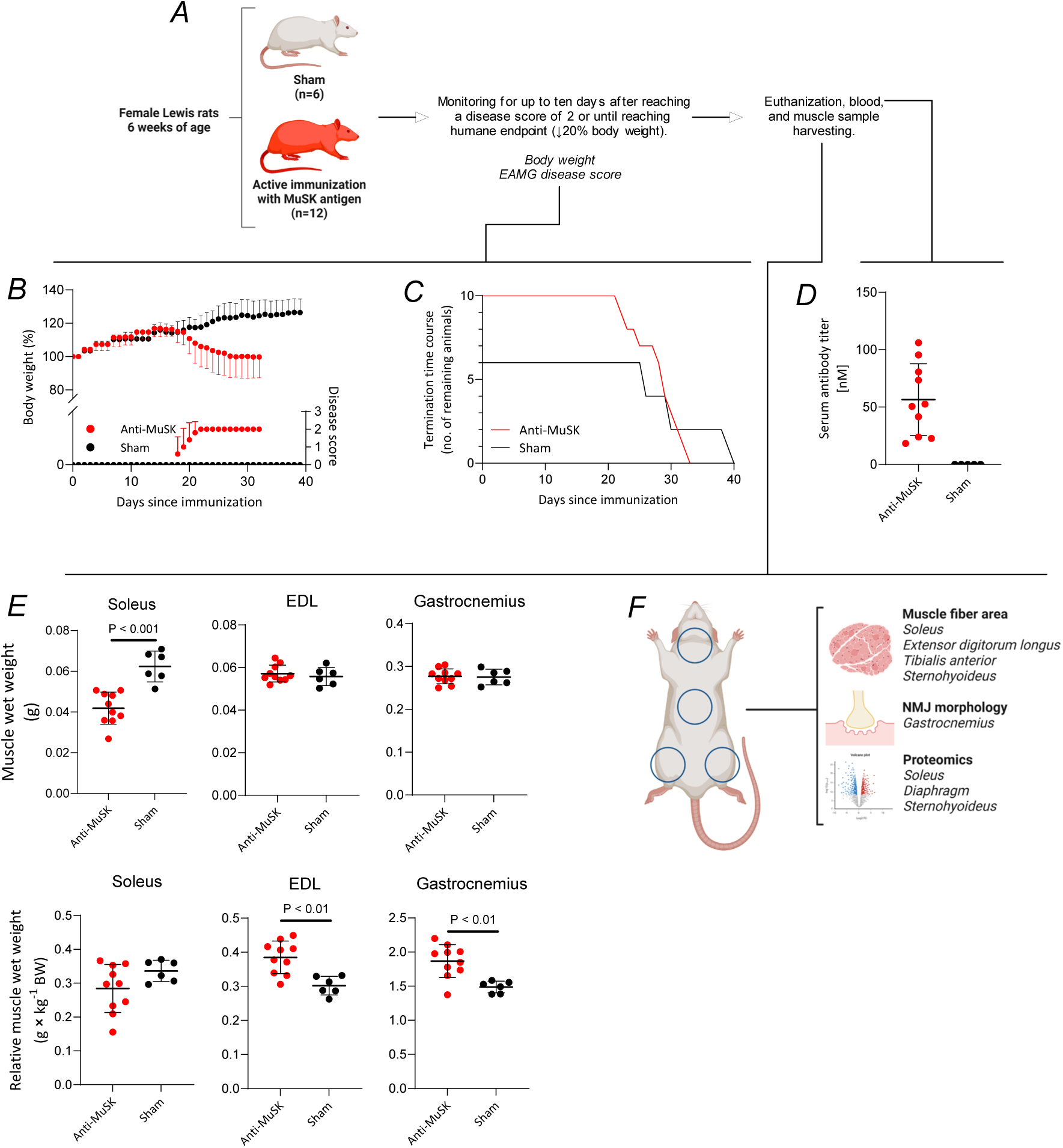
Study overview and animal characteristics. An overview of the study is shown in panel A. The development in body weight relative to the day of immunization, as well as the EAMG disease score, is shown in panel B and presented as the last observation carried forward. 10 out of 12 animals in the Anti-MuSK group developed disease symptoms and were thus included for further analyses. Panel C shows the number of remaining rats in each group from the time of immunization to the day the last rat was euthanized in the given group. While Anti-MuSK rats were terminated according to humane endpoints, or no later than 10 days after onset of a disease score of 2, Sham rats were terminated MuSK antibody concentration measured in serum of Anti-MuSK and Sham rats at the time of euthanization is shown in panel D. Panel E shows muscle wet weights of soleus, EDL, and gastrocnemius in either absolute numbers (upper row) or relative to body weight (lower row). Panel F depicts the analysis allocation of the harvested sample preparations of each muscle type. Lines (panel D, and E) and dots (panel B) are mean and error bars ± SD. P-values are based on an unpaired Student’s *T*-test.

### Animals

Female Lewis rats were purchased at 6 weeks of age from Janvier Labs (Le Genest-Saint-Isle, France) and enrolled at 7 weeks of age following a 1-week period of acclimatization. All rats were housed at 20-24 °C with a 12/12-hour light/dark cycle. Rats were fed with a standard diet and water *ad libitum*. 14 days after immunization, for animals undergoing active immunization (Anti-MuSK group), a soft (wetted) diet, as well as DietGel Recovery (cat. # 72-06-5022, ClearH2O), were supplied on the floor of the cage. Anesthetic procedures were performed by subcutaneous administration with fentanyl/fluanisol (Hypnorm®) and midazolam (Dormicum®) mixture. Euthanization was performed with an overdose of CO_2_. Humane endpoints for premature termination of rats were a ≥ 20 % decrease in body weight (compared to maximal measured body weight) and a disease score of ≥ 3 (see description below).

### Disease model

A previously established disease model by active immunization was used as the experimental animal model to induce a MuSK-MG disease phenotype (27,37). As antigen, the N-terminus peptide of a mouse MuSK 60 splice variant (N-MuSK60) was used for active immunization in combination with Complete Freund’s Adjuvant (CFA, cat. # 263810, BD Biosciences, Thermo Fisher Scientific) and Pertussis toxin (PT, cat. # P2980, Sigma-Aldrich). Rats were anesthetized and subcutaneously injected at the base of the tail with 10 µg N-MuSK60 peptide emulsified in CFA (150 µL total); 2.5 µg pertussis toxin was administered subcutaneously post-immunization. After the immunization procedure, a single injection of 2.5 µg PT was administered subcutaneously. The Sham-treated group received CFA and PT without antigen.

Afterward, rats in both groups were monitored under free-living conditions. Body weight and an MG-specific disease score were recorded as previously described (38) 3 times/week. In brief, the disease score was graded on a scale from 0 to 4 by the following criteria: no weakness (grade 0); fatigue or weakness after exercise (grade 1); a clinical sign of weakness before exercise, hunched back, head down (grade 2); severe clinical symptoms of weakness before exercise, no grip, hindlimb paralysis, compromised respiration (grade 3); moribund (grade 4). Rats were only included for further analysis if they reached a disease score of 2 within 30 days after immunization. Upon reaching a disease score of 2, rats were allowed to live for up to 10 additional days, with daily monitoring of body weight and disease scoring during this period. Sham rats were terminated in pairs within days of the Anti-MuSK counterparts (Figure 1C).

### Blood sampling and muscle tissue harvesting

At the time of termination, immediately before euthanasia, blood was collected from the tail vein for MuSK antibody titer measurements. Following euthanasia, muscle tissues were rapidly harvested and allocated for subsequent analyses. All tissues were rinsed in saline, blotted dry on paper, cleared of connective tissue, tendons, and residual blood, and weighed before freezing. From the left hindlimb, the soleus and extensor digitorum longus (EDL) muscles were excised, and from the thoracic cavity, the hemidiaphragm. These samples were snap-frozen in liquid nitrogen. From the right hindlimb, the whole gastrocnemius muscle and the mid-belly of the soleus, tibialis anterior (TA), and EDL were embedded in tissue freezing medium (Leica Biosystems, IL, USA) and frozen in dry ice–cooled isopentane. The sternohyoideus muscle from the neck region was longitudinally divided, with one half snap-frozen and the other embedded in tissue freezing medium at the mid-belly. All samples were stored at −70 °C until further analysis. For proteomic analysis, the soleus (hindlimb), diaphragm (respiratory muscle), and sternohyoideus (neck) were selected to represent distinct anatomical locations and muscle functions. The mixed-fiber type gastrocnemius muscle was examined to confirm the MG disease phenotype, characterized by denervated and fragmented endplates. Muscle atrophy was assessed by fiber type–specific morphological analysis in the hindlimb muscles (EDL, TA, soleus) and the sternohyoideus.

### MuSK antibody titer measurements by radioimmunoprecipitation assay (RIPA)

^125^I-labeling of the MuSK extracellular domain was performed using the chloramine T method, and antibodies in the test blood serum samples were quantified using RIPA. To summarize, 125i-MuSK, carrying 50,000 CPM, underwent a 2-hour incubation period with varying concentrations of the test serum, prepared in normal rat serum, at 4°C. The total rat serum employed was 2 μl. Subsequently, 10 μl of rabbit anti-rat serum was introduced, followed by an extended incubation period overnight at 4°C. Afterward, the samples were subjected to two wash cycles using 1 % Tween in PBS (PBST), and the residual radioactivity was quantified using a 1470 Wizard γ-counter (Perkin Elmer). Antibody titer (in [nM]) was calculated from the dilutions that exhibited a linear increase using the following formula:

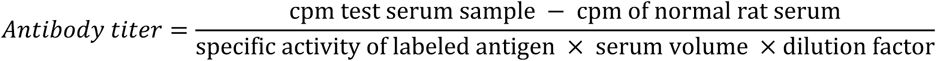

where the specific activity equals 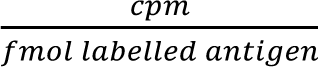

### Serum MuSK antibody staining in wild-type rat muscle

20 µm longitudinal sections of cryo-embedded soleus from a 14-week-old wildtype female Lewis rat were cut on a Leica CM3050 cryostat. For immunofluorescence of MuSK expression at the NMJ, slides were treated as described for NMJ morphology (see Immunohistochemistry NMJ morphology section below) using different antibodies. To further validate rat anti-MuSK seropositivity- and negativity of Anti-MuSK and Sham serum, respectively, serum from n=3 Sham and n=3 Anti-MuSK was diluted 1:100 in blocking buffer and used for overnight incubation of wild-type soleus slides at 4°C. Antibody titers in the selected Anti-MuSK animals were 106.08, 80.70, and 22.65 nM before dilution, and Sham titers were 0 nM. Secondary antibodies included donkey-anti-rat-488 (cat. # A48269TR, Invitrogen, 2 µg/mL) and α-BTX-555 to label rat-specific antibodies from serum at the NMJ. As positive control of MuSK protein localized at the NMJ, a staining using rabbit-anti-MuSK (cat. # ABS549, Sigma, 1:1000) and α-BTX-555 (α-BTX, cat. # B35451, Invitrogen, 4 µg/mL) + donkey-anti-rabbit-488 (cat. # 32790, Invitrogen, 2 µg/mL) was also performed. Sections were imaged on a Zeiss LSM800 laser scanning confocal microscope at the Bioimaging Core Facility, Health, Aarhus University, Denmark. To allow for comparison, the same imaging settings were used to acquire z-stacks of soleus incubated with serum from Sham and Anti-MuSK rats.

### Immunohistochemistry – muscle fiber morphology

To minimize potential variation introduced by disease period duration, only embedded muscle samples from Anti-MuSK rats terminated after 10 days with a disease score of 2 (i.e., non-humane endpoint termination, n = 5), as well as 5 Sham rats were included for immunohistochemical analyses. 10 µm transverse cryosections from embedded soleus, sternohyoideus, EDL, and TA muscle samples were prepared on microscope glass slides, air-dried, and stored at -80 °C until further analysis. A cryosection of one muscle type from an Anti-MuSK and a Sham rat were included on the same slide. In brief, following fixation with 4 % PFA, cryosections were blocked in 10 % normal goat serum (cat. # ab7481, Abcam) in PBS at room temperature. Primary antibody staining was performed for 2 h in 10 % goat serum in PBS at room temperature. The primary antibodies were used to stain muscle fiber borders (laminin polyclonal rabbit IgG antibody, 1:200 dilution, cat. # PA1-16730, Invitrogen, Thermo Fisher Scientific) and myosin heavy chain I (MHC-I) positive fibers (MYH7 monoclonal mouse IgG2b antibody, 1:50 dilution, cat. # BA-F8, Developmental Studies Hybridoma Bank, DSHB, developed by Schiaffino, S). Secondary antibody staining was performed in darkness for 1 h at room temperature using goat anti-rabbit IgG Alexa fluor 647 (1:250 dilution, cat. # A-21245, Invitrogen) and goat anti-mouse IgG2b (Alexa fluor 488, cat. # A-21141, 1:500 dilution, Invitrogen). Finally, coverslips were mounted using a mounting medium (Invitrogen ProLong™ Gold Antifade Mountant, Cat. # P10144, Thermo Fisher). Glass slides were then dried and stored at 4 °C until fluorescence microscopy analysis.

### Immunohistochemistry – NMJ morphology

For the NMJ morphology analyses, muscle preparations from the same rats used for the muscle fiber morphology analyses were selected (n = 5 per group). 20 µm longitudinal sections of embedded gastrocnemius were cut on a Leica CM3050 cryostat, air-dried, and stored at -70°C until further use. For immunofluorescence of the NMJ, slides were fixated for 10 min with 4% paraformaldehyde, permeabilized with PBS + 0.1% Triton X-100, before being blocked for 30 min with 10% donkey serum (cat. # D9663, Sigma) in PBS + 0.1% Triton -X-100 at room temperature. Primary antibody staining was performed overnight at 4°C in blocking buffer with mouse anti-rat neurofilament M (NF-M, cat. # 2H3, DSHB, developed by Jessell, T.M. / Dodd, J, 0.58 µg/mL concentration applied) and mouse-anti-synaptic vesicle glycoprotein 2 (cat. # SV2, DHSB, developed by Buckley, K.M., 0.55 µg/mL concentration applied) to label nerve. The following day, sections were washed with PBS + 0.1% Triton X-100 before being incubated with secondary antibodies in darkness for 3 h at room temperature using α-Bungarotoxin-555 to label AChRs (α-BTX, cat. # B35451, Invitrogen, 4 µg/mL) + donkey-anti-mouse-488 (cat. # A32766, Invitrogen, 2 µg/mL). Sections were washed with PBS followed by Hoechst in PBS (cat. # 33342, Thermo Fisher Scientific, 2 µg/mL) for 10 min to label nuclei, before being mounted with Dako fluorescent anti-fading mounting medium (cat. # S302380, Agilent). Glass slides were then dried and stored at 4 °C until imaging.

### Muscle fiber morphology analysis

Images for muscle fiber morphology were obtained by fluorescence microscopy using an EVOS M7000 automated imaging system (Invitrogen). Whole muscle cross-sections were scanned and captured at 20x magnification. All sections were assessed for eligible fibers, meaning nonorthogonal cut fibers and fibers with poor sarcolemmal and morphological definition were excluded. A semiautomatic approach was applied to obtain fiber area and minimum ferret diameter measurements through the MATLAB (The MathWorks Inc.) plugin SMASH (39). To perform fiber type-specific quantification in slow-twitch muscle fibers, type I fibers were identified as MHC-I (MYH7) positive fibers. Given the vast dominance of type I slow-twitch fibers in soleus muscle of rodents, fiber type analyses were not performed in this muscle type. For soleus, EDL, and sternohyoideus, all eligible fibers on all sections were quantified. For TA, 5-6 representative sections were chosen in which all eligible fibers were quantified. On average, 1837 ± 383 fibers (mean ± SD) were quantified per animal in soleus. In EDL, an average of 1901 ± 365 fibers were quantified per animal, of which 58 ± 37 were identified as type I. In TA, an average of 2562 ± 270 fibers were quantified per animal, of which 45 ± 13 were identified as type I. In sternohyoideus, an average of 2092 ± 823 fibers were quantified per animal, of which 48 ± 34 were identified as type I. The investigator performing the quantification was blinded against the groups (Anti-MuSK and Sham) through random ID generation of image file names.

### NMJ morphology analysis

A Zeiss LSM800 laser scanning confocal microscope with a 40X oil objective (Bioimaging Core Facility, Health, Aarhus University, Denmark) was used to acquire z-stacks projections of en-face NMJs. Images were analyzed using ImageJ software where endplates were manually traced according to AChR signal and AChR and nerve area measured. The number and size of AChR fragments were determined using “Analyze Particles” with a size from 1 µm^2^-infinity. Co-localization of AChR and nerve was calculated using the freely available plugin “Colocalization” and a ratio of 50%. Innervation degree was scored as innervated (pre-synapse signal of SV2 and NF-M occupies >90% of post-synapse AChR signal), partially innervated (pre-synapse vacates >10% of post-synapse), or denervated (pre-synapse occupies <10% of post-synapse). 5 animals in each group were analyzed with 15.4 ± 0.9 NMJs in the Sham group and 17.2 ± 1.5 NMJs in the Anti-MuSK group (mean ± SD). A mean for each animal was calculated and used as data point for the statistical analyses.

### Protein extraction and preparation for LC-MS/MS

Snap-frozen soleus, diaphragm, and sternohyoideus muscle tissues were pulverized in a liquid nitrogen-cooled mortar with a chilled pestle and transferred to protein LoBind® tubes (Eppendorf). Due to limited tissue availability, one Anti-MuSK diaphragm replicate was excluded. Muscle peptide digests for LC-MS/MS analysis were prepared using a filter-aided sample preparation (FASP) protocol (40,41). Tissue lysis was performed in 4% SDS and 100 mM Tris/HCl (pH 7.6), freshly supplemented with 100 mM DTT, using 10 µl buffer per mg tissue. Samples were vortexed, heated at 95°C for 5 min, sonicated for 2 × 20 s (Branson probe), and centrifuged at 16,000 × g for 10 min to remove debris. Protein concentration was determined using the Pierce™ 660 nm protein assay kit with detergent compatibility reagent (cat. #22660 and #22663, Thermo Fisher Scientific, Waltham, MS, USA).

For FASP, 100 µg protein was combined with 200 µl of buffer UA (8 M urea, 100 mM Tris/HCl, pH 8.5) in pre-conditioned 30 kDa centrifugal filter units (Amicon Ultra-0.5, cat. #UFC5030, Millipore). After centrifugation (10,000 × g, 20°C, 15 min), two more washes with 100 µl of buffer UA were performed. Alkylation was achieved by adding 50 µM iodoacetamide in 100 µl buffer UA, mixing (600 RPM for 1 min), and incubating for 20 min in the dark at room temperature. After two additional washes with buffer UA, the filters were washed twice with 100 µl of buffer UB (4 M urea, 100 mM Tris/HCl, pH 8.0). Protein digestion was initiated by adding Lys-C (cat. # VA1170, Promega, 1:50 enzyme-to-protein ratio) in 40 µl buffer UB and incubating at 37°C for 1 h with gentle mixing. Trypsin (cat. # V5280, Promega, 1:100 enzyme-to-protein ratio) was then added in 120 µl of 50 mM ammonium bicarbonate (buffer ABC), and digestion proceeded for 16 h at 37°C. Peptides were recovered through three centrifugation steps (10,000 × g, 20°C, 10 min) with 100 µl of buffer ABC added between each. Enzyme activity was quenched with 3% trifluoroacetic acid, and peptide digests were concentrated in a SpeedVac centrifuge followed by cleaning with C18 spin columns (cat. # 89873, Thermo Fisher Scientific). TMT labeling was performed using 16plex tandem mass tag reagents (TMTpro™ 16plex, cat. # A44522, Thermo Fisher Scientific) according to the manufacturer’s protocol. Three TMT plexes (soleus, diaphragm, and sternohyoideus) were constructed to compare the Anti-MuSK and Sham muscle proteomes. After quenching, TMT-labeled samples were pooled, cleaned with strong cation exchange (SCX) cartridges (Strata® SCX, cat. # 8B-S010-EAK, Phenomenex) and concentrated in a SpeedVac centrifuge. Peptides were fractionated using isoelectric focusing (IEF) with immobilized pH gradient strips (Cytiva Immobiline™ DryStrip, pH 3-10, linear gradient, 18 cm, cat. # GE17-1235-01, Cytiva Life Sciences). Concentrated peptides were reconstituted in IEF buffer (350 µl) and mixed with rehydration buffer (450 µl) before loading into rehydration lanes. Gel strips were rehydrated for 15 h under 1 ml of paraffin oil and subjected to electrophoresis in four steps (500 V for 1 min, 3500 V for 1.5 h, 3500 V for 12 h, and 800 V for 12 h). Gel strips were cut into 16 fractions, and peptides were extracted with an extraction buffer (5% acetonitrile, 0.5% trifluoroacetic acid) for 1 h under gentle shaking. Peptide fractions were then purified using Pierce™ C18 spin columns (cat. # 89870, Thermo Fisher Scientific), according to the manufacturer’s protocol, and concentrated in a SpeedVac centrifuge before LC-MS/MS analysis.

### LC-MS/MS-based proteomics

The peptide mixtures were analyzed by nano-liquid chromatography tandem mass spectrometry (nanoLC-MS/MS) (EASY nanoLC-1200, Thermo Fisher Scientific) coupled to Q Exactive™ HF-X Quadrupole-Orbitrap™ Mass Spectrometer (Thermo Fisher Scientific) as previously described (42). TMT labeled peptide samples were trapped on a pre-column (PepMap 100, 2 cm, 75 μm i.d., 3 μm C18 particles, 100 Å, Thermo Scientific) followed by separation on a C18 column with integrated emitter (EASY-Spray column, PepMap 25 cm, 75 μm i.d., 2 μm, 100 Å, Thermo Scientific). Peptides were separated in a 100 min linear gradient, from 4 to 40% acetonitrile in 0.1% formic acid at a flowrate of 300 nL/min. The MS was operated with data dependent acquisition in positive mode, automatically switching between precursor scanning (MS1) and fragmentation (MS2) acquisition. Resolution of MS1 was set to 60,000 and MS2 to 45,000. In MS1, automatic gain control target was set to 1 × 10^6^ ions and scan range between 372 and 1500 m/z. In MS2, AGC target was set at 2 × 10^5^ ions, with fixed first mass set to 120 m/z. Dynamic exclusion was set to 20 s for all analyses. Up to twelve of the most intense ions were fragmented per every full MS scan, by higher-energy C-trap dissociation at the setting of 35. Ions with single charge or unassigned charge states were excluded from fragmentation.

### Proteomics data analysis and bioinformatics

Database searches of the raw MS peptide spectra were performed in Proteome Discoverer v 3.0 (Thermo Fisher Scientific) using a Uniprot database containing 8,284 *Rattus norvegicus* sequences (downloaded Oct 29^th^ 2023). At least 1 unique peptide per protein was required for protein identification, and up to two missed trypsin cleavages were allowed. Lists of protein groups with abundance ratios from each TMT-plex analysis were exported from Proteome Discoverer. Only protein groups identified with high confidence (false discovery rate, FDR, < 0.01) were included for analysis. No imputation of missing data was performed, meaning that protein groups with a missing signal in at least one TMT channel were excluded. A Sham sternohyoideus replicate from TMT study 3 was excluded from data analysis due to a manual handling error in the lab during a SpeedVac step. Raw protein abundance ratios were log_2_ transformed followed by normalization by median subtraction of the overall sample median, and scaling of the individual protein abundance against the median across samples (43). The DAVID tool (44,45) was used to annotate data sets with Gene Ontology (GO) terms, and additionally to perform enrichment analysis of the identified proteins against the rat background genome (GO cellular components, Figure 4B). For hierarchical clustering and visualization (Figure 4D), data was imported into the Perseus software (version 2.0.1.1 (46)) and analyzed with default settings. For analysis of differentially expressed proteins (Figure 5A), abundance ratios were compared between Anti-MuSK versus Sham within each muscle type/TMT-plex using the *limma*-based [S9] *R*-package *DEqMS* [S9] for multiple testing with FDR-correction. Afterward, proteins were ranked based on a fusion significance score (π-value (47)) calculated as

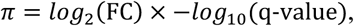

where log_2_(FC) is the fold-change calculated as the difference between the log_2_ means of Anti-MuSK versus Sham, and *q*-value is the FDR-corrected *P*-value from *DEqMS* results. These calculations were performed in *Rstudio* (RStudio Team, Boston, MA) with related visualizations performed with the *R*-package *ggplot2* (48).

For gene set enrichment analysis using *GSEA* (version 4.3.3 (49,50)), lists of proteins for each differential expression analysis were prepared, ranked by the π-value, and annotated with the corresponding gene symbol. GSEA was performed separately for selected GO and curated gene sets based on .gmt files (GOCC, GOBP, Reactome, and WikiPathways) downloaded from the *MSigDB* database (51–53) (version 2023.2). GSEA parameters were set to consider only terms with ≥5 and ≤300 genes and executed with 10^5^ permutations. Afterward, *P*-values from the GSEA output were FDR-corrected (to *q*-values), per the Benjamini-Hochberg procedure, using the *R*-package *qvalue* (54). Gene sets with FDR *q*-value <0.05 were considered significantly enriched in either up- or downregulated direction based on the enrichment score (ES). For network visualization of GO-terms (Figure 6 and Supplemental Figure S2), GSEA results were imported into *Cytoscape* (version 3.10.1 [S20]) using the *EnrichmentMap* plugin (version 3.3.6 (55)), with the addition of gene set cluster analysis visualization using the *AutoAnnotate* plugin (version 1.4.1 (56)) executed with default MCL-cluster settings. Annotations based on the *MitoCarta 3.0* catalog (57) were used for in-depth analysis of the identified mitochondrial proteome (Figure 7).

### Other statistical analyses

For the comparisons of muscle weights (Figure 1), NMJ morphology (Figure 2), and muscle fiber area (Figure 3) between Anti-MuSK and Sham animals, the unpaired student’s *t*-test was used. For analysis of the innervation profile (Figure 2D), a two-way ANOVA was applied with groups (Anti-MuSK and Sham) and innervation (fully, partially, and denervated) as the main effects. α-value was set to 0.05. All values are reported as mean ± SD, unless otherwise specified.

**Figure 2.**
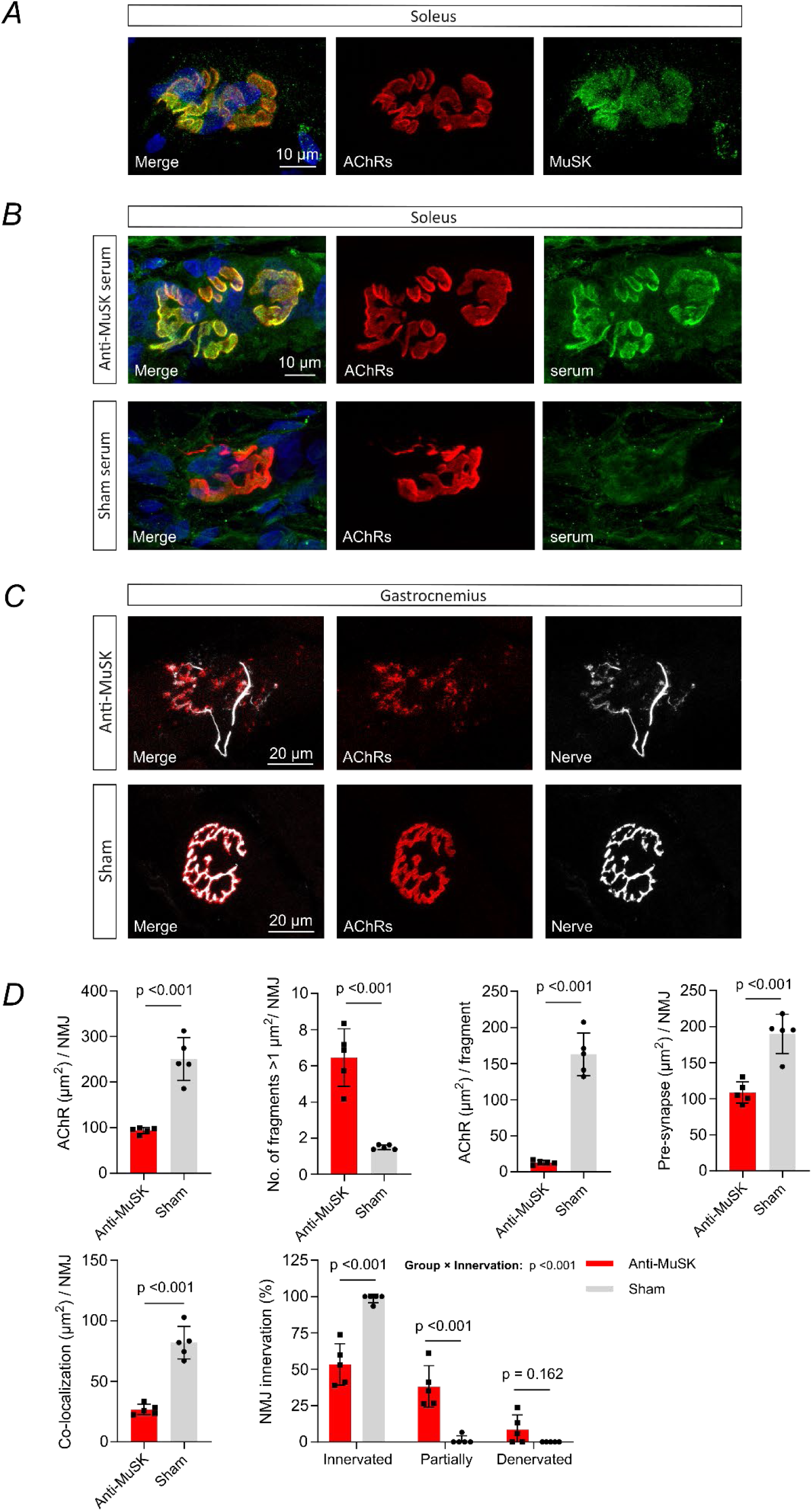
Immunohistochemical analyses of serum MuSK antibody reactivity and neuromuscular junction (NMJ) morphology. Panel A depicts representative confocal microscopy images of immunofluorescence staining of an NMJ/endplate as visualized by the acetylcholine receptors (AChRs, α-bungarotoxin staining) and MuSK protein in longitudinal cryosections from wild-type soleus muscle. The post-synaptic co-localization and clustering of MuSK and AChR in wild-type muscle was thus confirmed. Panel B shows representative images confirming the localization of MuSK antibodies at the post-synaptic area of the NMJ in longitudinal cryosections from wild-type soleus muscles following incubation with serum obtained from Anti-MuSK rats (upper panel row). The lower row in panel B confirms seronegativity in Sham serum as indicated by only the presence of background signal from the Rat IgG secondary antibody. For representative images of staining from incubation of all the selected serum samples (n = 3 from group), see the supplemental figure S1. Panel C shows representative images of an identified NMJ by colocalization of post-synaptic AChRs and pre-synaptic nerve terminal markers synaptic vesicle protein 2A (SV2) and neurofilament M protein (NF-M) in gastrocnemius muscle from Anti-MuSK (upper row) and Sham (lower row) rats. The observations of fragmented and scattered AChR clusters in Anti-MuSK gastrocnemius visually confirm the myasthenia gravis phenotype. Panel D shows results from the analysis of NMJ morphology in gastrocnemius muscles from Anti-MuSK and sham rats, with mean AChR area, mean number of fragments, mean AChR area per fragment, pre-synaptic area, co-localization of pre- (SV2+NF-M) and post-synaptic (AChRs) markers, and the NMJ innervation profile. 15.4 ± 0.9 NMJs in the Sham group and 17.2 ± 1.5 NMJs in the Anti-MuSK group (mean ± SD) were analyzed. Bars are mean ± SD, and the numbers above bars represent *P*-values derived from the student’s unpaired *t*-test.

## 3. Results

### Disease phenotype in active immunized rats

Following the immunization procedure, a disease phenotype (as indicated by a disease score of 2) was observed within 17 to 21 days after immunization (Figure 1B). Two rats in the Anti-MuSK group did not develop an adequate disease phenotype based on the prescribed criteria and were thus excluded from the study (see *Disease model* methods section). While Sham rats exhibited an expected weight gain throughout the study period, Anti-MuSK rats lost an average of 12.5 ± 6.6% body weight from the onset of disease symptoms to the day of euthanization (Figure 1B). On average, the Anti-MuSK rats were terminated 26.4 ± 3.8 days after immunization (Figure 1C), with n = 5 terminated due to humane endpoint (HE, loss of body weight), and n = 5 following ten days living with a disease score of 2 (non-HE). Sham rats (n = 6) displayed no disease symptoms throughout the study period and were terminated at an average of 30.8 ± 6.2 days after sham-immunization to accommodate the main study purpose. Circulating MuSK antibodies were present in all included Anti-MuSK rats, whereas Sham rats were seronegative (Figure 1D).

The wet weight of the left hindlimb slow-twitch dominated soleus muscle measured immediately following euthanization was approximately 32% less in Anti-MuSK rats than that of Sham (0.042 ± 0.008 g versus 0.062 ± 0.008 g, *P* < 0.001), whereas the absolute wet weights of EDL and gastrocnemius muscles were comparable between groups (Figure 1E). Normalizing muscle wet weights relative to body weight eliminated the difference in soleus muscle weight between groups, whereas the relative weights of EDL and gastrocnemius muscles were approximately 27% and 26% greater, respectively, in Anti-MuSK rats compared to Sham (*P* < 0.01 for both comparisons).

To further confirm the presence of MuSK-specific antibodies in serum derived from Anti-MuSK rats, immunohistochemical analysis with serum incubation in wild-type rat soleus muscle was performed using serum from Anti-MuSK and Sham rats (n = 3 each group, Figure 2B and Supplemental Figure S1). The three seropositive Anti-MuSK animals were selected to represent the full range of measured ab titer values. The analysis confirmed post-synaptic co-localization of IgG-antibodies and AChRs in muscle sections incubated with Anti-MuSK serum, as opposed to the presence of only background signal when incubated with Sham serum.

### NMJ morphology

The impact of active immunization with MuSK-antigen on NMJ morphology was evaluated in gastrocnemius muscle by immunohistochemical analysis using confocal microscopy. By staining pre-synaptic nerve terminals (using SV2 and NF-M antibodies) and post-synaptic AChR clusters (using α-bungarotoxin), a clear disease phenotype was observed in Anti-MuSK muscles (Figures 2C and 2D and Supplemental Figure S1). Loss of AChR area (*P* < 0.001), as well as increased fragmentation of the NMJ (*P* < 0.001), was observed in Anti-MuSK skeletal muscle as opposed to those of the Sham group (Figure 2D). Loss of pre-synaptic area was likewise evident in Anti-MuSK muscles (Figure 2D, *P* < 0.001). Moreover, while the innervation profile of Sham muscles was characterized as fully innervated (Figure 2D), an average of 9 ± 10 % of the identified NMJs were characterized as denervated, 38 ± 14 % partially innervated, and 53 ± 14 % innervated in Anti-MuSK muscles.

### Muscle fiber morphology

Immunohistochemical analysis (Figure 3) revealed marked muscle fiber atrophy in the slow-twitch/type I fiber-dominant soleus muscle of Anti-MuSK rats as compared to Sham (*P* < 0.001). Accordingly, mean min. Feret diameter of soleus muscle fibers were 33.0 ± 3.3 µm and 43.5 ± 1.9 µm in Anti-MuSK and Sham animals, respectively. While overall min. Feret muscle fiber diameter was similar between groups in EDL, tibialis anterior, and sternohyoideus muscles (Figure 3B-D), the diameter of identified MYH7^+^-fibers (type I, slow-twitch) was smaller in Anti-MuSK rats as compared to Sham. In EDL muscles, mean Feret diameter of type I fibers was 18.9 ± 1.0 µm and 23.3 ± 2.1 µm (*P* < 0.01), in tibialis anterior 18.4 ± 0.8 µm and 28.9 ± 2.3 µm (*P* < 0.001), and in sternohyoideus 16.4 ± 1.5 µm and 22.0 ± 0.8 µm (*P* < 0.001), for Anti-MuSK and Sham rats, respectively. Finally, no impact on type II min. Feret fiber diameter was observed in Anti-MuSK.

**Figure 3.**
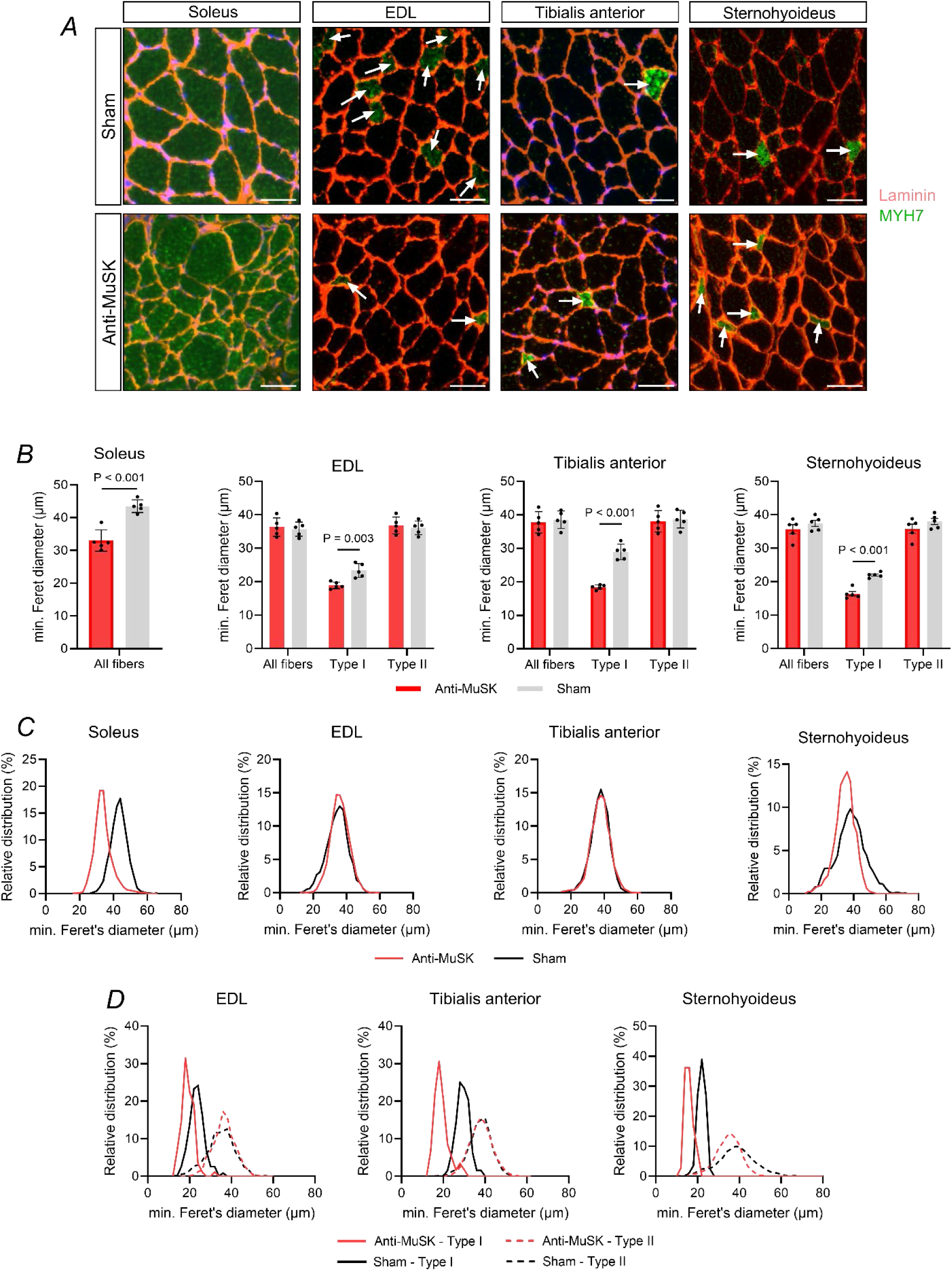
Immunohistochemical analyses of muscle fiber size in skeletal muscles. Panel A depicts representative images of immunofluorescence staining in transverse cryosections from soleus, EDL, tibialis anterior, and sternohyoideus muscles of Sham and Anti-MuSK rats. White arrows mark identified type I fibers (MYH7^+^) in EDL, tibialis anterior, and sternohyoideus muscles. Panel B shows min. Feret’s diameter of all muscle fibers in the type I-dominant soleus muscle and for type I and II fibers in EDL, tibialis anterior, and sternohyoideus. The corresponding relative distribution of min. Feret’s diameter of all fibers is shown in panel C, and the fiber-type specific relative distribution in panel D. Bars are mean ± SD, and the numbers above bars represent *P*-values derived from the student’s unpaired *t*-test. The white scale bar is 50 µm.

### Identified proteins by whole-muscle proteomics

Across both Anti-MuSK and Sham, the proteomics analysis workflow identified 1868, 2135, and 2046 proteins with high confidence (FDR <0.01) in soleus, diaphragm, and sternohyoideus muscles, respectively (Figure 4A). 1574 of the identified proteins was shared between the three muscle types (Figure 4A), in which mitochondrial, ribosomal, and cytoplasmic terms were among the top enriched cellular components, as revealed by enrichment analysis of GOCC terms against the rat genome background (Figure 4B). Among manually selected muscle-relevant GOCC terms, sarcoplasmic reticulum, Z-disc, T-tubuli, and sarcomere cellular components exhibited the highest coverage of identified proteins in all three muscle types (Figure 4C).

**Figure 4.**
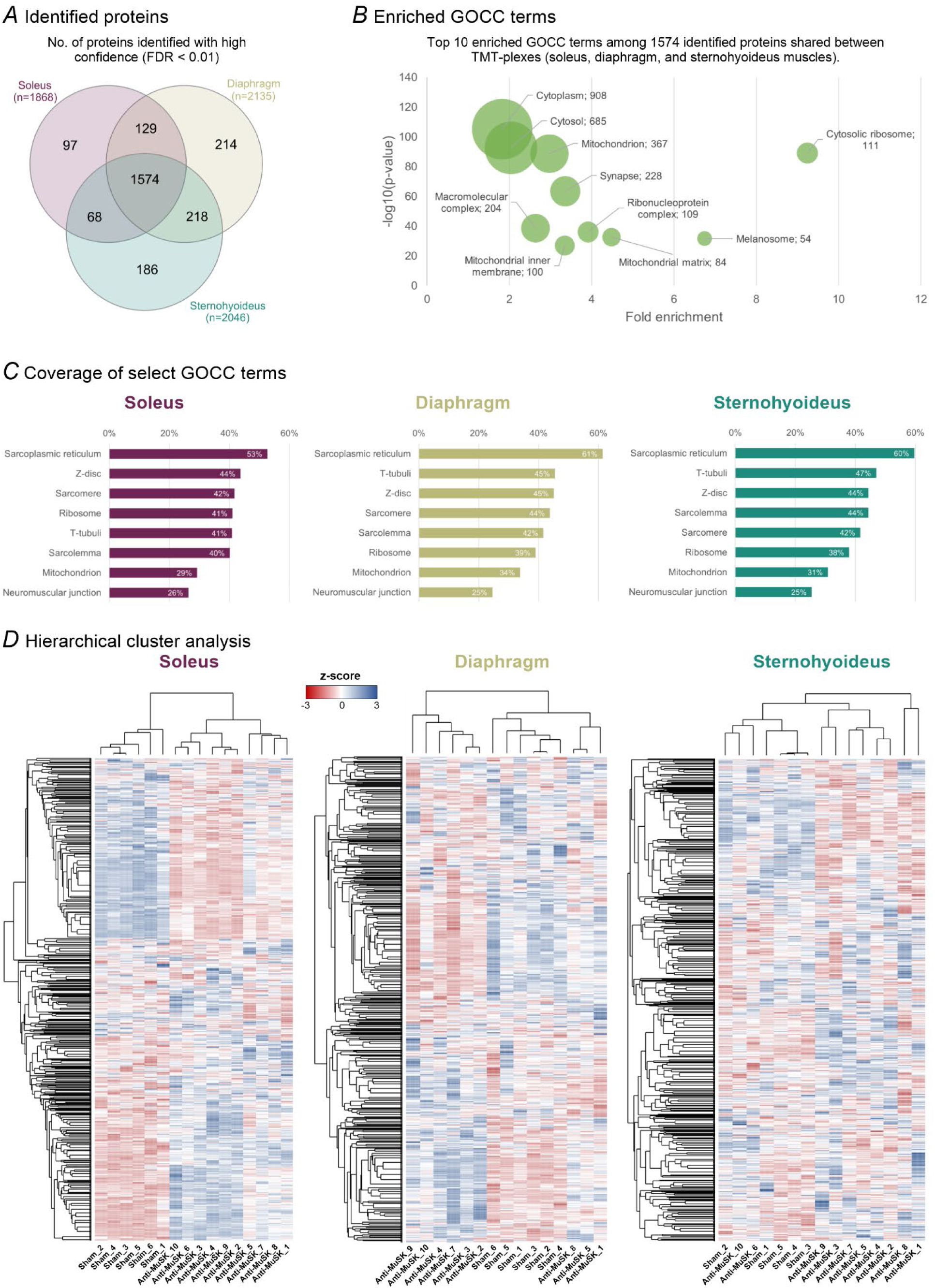
LC-MS/MS-based bottom-up proteomics performed in whole-muscle protein extracts from three different muscle types. The Venn diagram in Panel A illustrates the overlap in identified proteins with high confidence (FDR < 0.01) between the three analyzed muscle types. Panel B depicts enriched gene set terms (FDR < 0.05) from the gene ontology cellular components (GOCC) category among the identified 1574 proteins shared between the three muscle types, by comparing proteins against the rat genome as background using the DAVID functional annotation tool. Numbers denote the number of identified proteins within the given gene set term. The horizontal bar graph in panel C shows the coverage of select muscle-related GOCC terms (number of identified proteins versus the gene set term total in percentage). Panel D visualizes unbiased hierarchical cluster analysis conducted using the Perseus software for each TMT-plex/muscle type run.

Hierarchical cluster analysis (Figure 4D) revealed two overall column clusters separating the Anti-MuSK and Sham soleus muscle proteomes, with two further distinct column sub-clusters among Anti-MuSK samples. For the diaphragm, two overall column clusters were also identified, with an Anti-MuSK only cluster formed of six samples, whereas the other column cluster included all Sham samples as well as the remaining three Anti-MuSK samples separated into two sub-clusters. The proteome of sternohyoideus did not exhibit a distinct column clustering between Anti-MuSK and Sham.

### Differentially expressed proteins (DEPs) in Anti-MuSK versus Sham skeletal muscle

Comparing protein abundances between Anti-MuSK and Sham phenotypes, using the *limma*-based *DEqMS R*-package, revealed a muscle type-dependent impact of the MuSK active immunization (Figure 5A). Accordingly, the proteome of Anti-MuSK soleus muscles was impacted to the greatest extent, with the abundance of 621 and 540 proteins being differentially expressed in either up- or downregulated direction, respectively, in comparison to Sham (q <0.05). For diaphragm muscle, 368 and 376 proteins were up- and downregulated, while the sternohyoideus muscle type was affected the least in the Anti-MuSK phenotype with 99 and 109 up- and downregulated proteins, respectively (q <0.05). Results of the DEP analysis are available as supplementary data (supplementary table S4-6). The number of overlapping up- and downregulated proteins between muscle types is depicted in Venn diagrams of Figure 5B.

**Figure 5.**
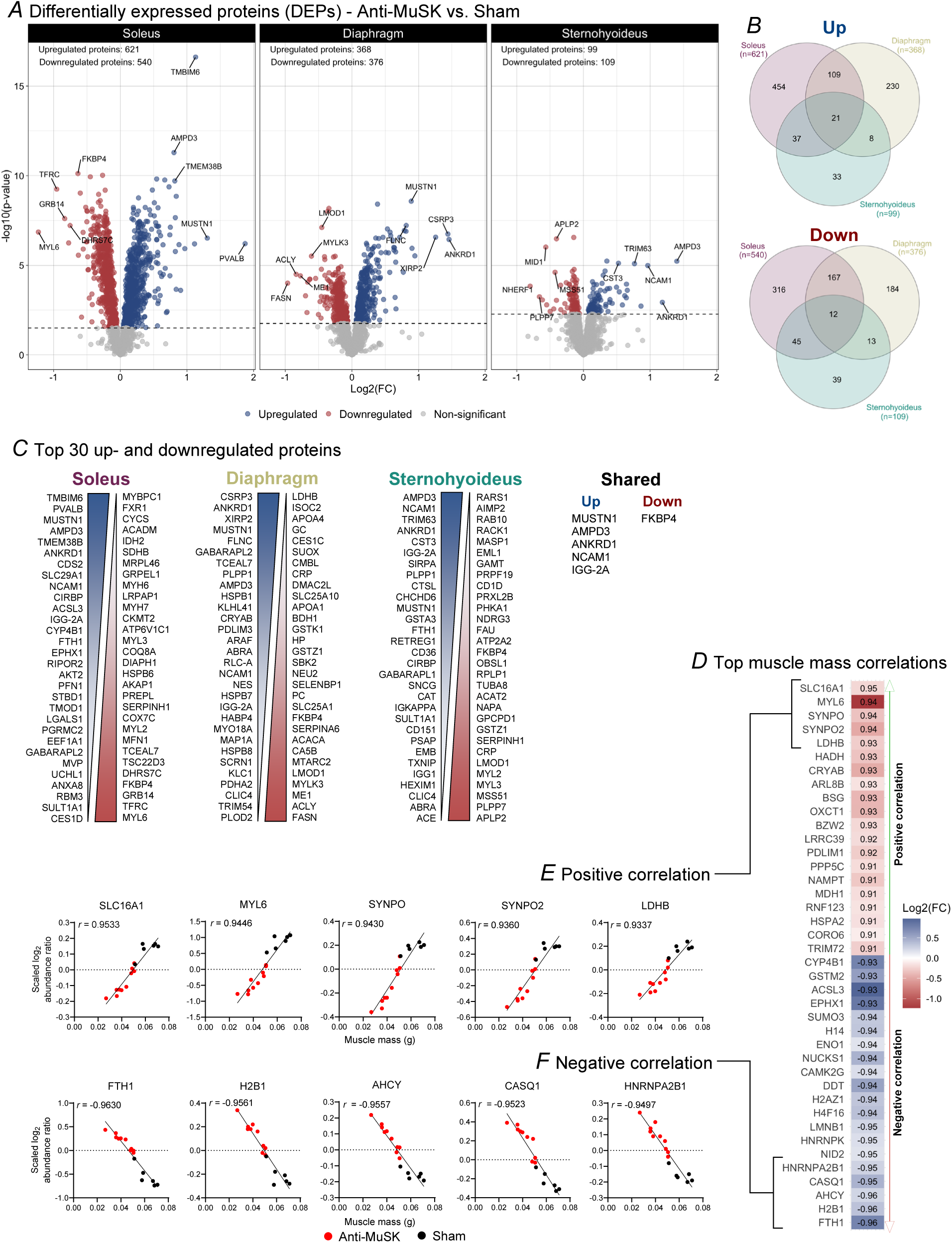
Differential expression analysis in Anti-MuSK versus Sham rats and associations between soleus muscle mass and protein abundance. Volcano plots visualizing the results of the differential expression analysis, - log_10_(*P*-value) plotted against the log2FC (Anti-MuSK versus Sham) conducted within each muscle type using the *limma*-based *R*-package *DEqMS* are shown in panel A. Dashed lines depict the significance cut-off (FDR q-value <0.05), with proteins identified as differentially expressed in either an up- (log2FC >0) or downregulated (log2FC <0) direction. Based on the π-value significance score, the top 5 regulated proteins in both directions within each muscle type analysis are highlighted. Venn diagrams in panel B represent the number of regulated proteins in up- and downregulated direction, respectively, within and between each muscle type in the Anti-MuSK phenotype compared to Sham. Panel C depicts the top 30 regulated proteins in both directions within each muscle type and shared between them in the Anti-MuSK phenotype based on ranking by π-value. The top 20 positive and negative correlations between soleus muscle mass and protein abundance are displayed in panel D. The numbers within each tile represent the Pearson correlation (*r*) and color represents the log2 fold change (Anti-MuSK versus Sham) as indicated by the scale bar. Panels E and F highlight the individual data points of protein abundance plotted against soleus muscle mass of the top 5 positive and negative correlations (based on *r*), respectively.

Ranking the top 30 differentially expressed proteins in both directions by π-value (Figure 5C) revealed an overlap in the top-regulated proteins, with MUSTN1, AMPD3, ANKRD1, NCAM1, and IGG-2A being upregulated, and FKBP4 downregulated in Anti-MuSK versus Sham across all three muscle types.

### Associations between soleus muscle mass and protein abundance

Pearson correlation coefficients (*r*) were calculated to assess the linear relationship between soleus muscle mass and the individual abundance of all identified proteins in the soleus muscle (Supplementary Table S7). Figure 5D displays the proteins with the highest 20 positive and negative correlation coefficients, while Figures 5E and 5F plot the top 5 coefficients, showing protein abundance against muscle mass. Notably, strong negative correlations (*r* = -0.96 to -0.95) were observed between soleus muscle mass and the abundance of FTH1, H2B1, AHCY, CASQ1, and HNRNPA2B1. Conversely, strong positive correlations (*r* = 0.93 to 0.95) were found between soleus muscle mass and the abundance of LDHB, SYNPO2, SYNPO, MYL6, and SLC16A1.

### Gene set enrichment analysis

Computational bioinformatics analysis was conducted using gene set enrichment analysis (GSEA) to explore regulatory themes in Anti-MuSK skeletal muscle. We utilized curated gene sets from the *MSigDB* database, including cellular components (*GOCC*), biological processes (*GOBP*), and pathways (*Reactome* and *WikiPathways*) [S15-S17], to examine the enrichment of regulatory themes among the up- and downregulated proteins in Anti-MuSK skeletal muscle across different muscle types. For GSEA, the significance score (π) was employed to rank the regulation of all identified proteins within each muscle type. Full GSEA results are provided in supplementary data (Supplementary Tables S8-10).

Figure 6 illustrates networks of enriched cellular components (FDR q < 0.05) among regulated proteins within muscle types and shared terms between them. The diaphragm muscle prominently features enriched cellular components in the upregulated direction (enrichment score, *ES* > 0, Figure 6A). Notably, protein complexes such as the proteasome and peptidase complex, cytoskeletal components like the sarcomere, microtubules, and contractile fibers as well as ribosomal components including ribosomal subunits and the ribonucleoprotein complex, form distinct network clusters of upregulated components in the Anti-MuSK diaphragm muscle. In contrast, no such network clusters of upregulated cellular components were observed in the Anti-MuSK soleus or sternohyoideus muscles.

**Figure 6.**
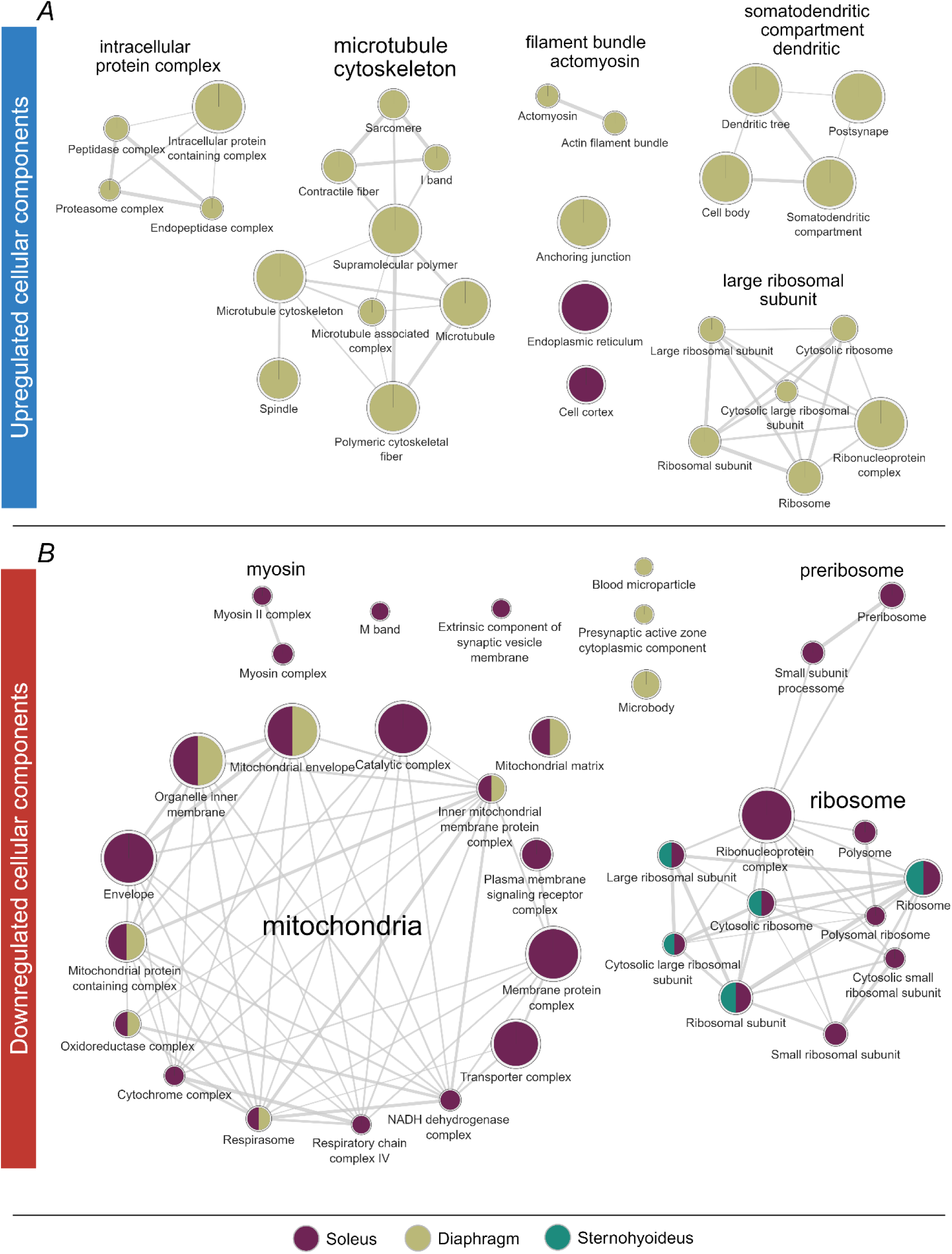
Networks of regulated cellular components in Anti-MuSK skeletal muscles. Network visualization of enriched GOCC terms in the up- (enrichment score >0) and downregulated (enrichment score <0) direction are displayed in panels A and B, respectively. Only significantly enriched terms (FDR q <0.05) as evaluated by gene set enrichment analysis were used for visualization. Networks were created using *Cytoscape* and the plugins of *EnrichmentMap* and *AutoAnnotate* by importing the results of the gene set enrichment analyses from pre-ranked proteins lists (based on π-value) on curated GOCC gene sets. The headline of each node network represents a broader annotation of the gene terms in the specific group. Each node represents a GOCC term, with node size visualizing the number of proteins in that term. Edge thickness between nodes represents the fraction of shared proteins between terms. Single nodes without edges represent significantly enriched terms with no shared proteins in other terms.

In the downregulated direction (*ES* < 0), two distinct network clusters emerged, representing mitochondrial and ribosomal components (Figure 6B). The mitochondrial cluster included cellular components such as respiratory chain protein complexes (e.g., NADH dehydrogenase, oxidoreductase, complex IV, and cytochrome complex) and mitochondrial structures (e.g., inner membrane and envelope). The soleus muscle was consistently represented across all enriched terms within this cluster, while the diaphragm muscle showed an overlap in some downregulated terms. The ribosomal cluster comprised enriched terms related to ribosomal subunits, the ribonucleoprotein complex, and pre-ribosomal components, found either in the soleus and sternohyoideus muscles or in the soleus alone. Additionally, a smaller cluster related to downregulated myosin complexes was identified exclusively in the soleus muscle of Anti-MuSK samples.

Network visualizations of regulated biological processes and bar graphs of enriched Reactome and WikiPathways terms are provided in the supplementary figures (Supplementary Figures S2 and S3).

### Skeletal muscle mitochondrial deficits in Anti-MuSK rats

Protein annotation using the MitoCarta 3.0 catalog further highlighted the significant impact of MuSK active immunization on the skeletal muscle mitochondrial proteome (Figure 7). A distinct, muscle type-dependent effect was evident in Anti-MuSK muscles (Figure 7A). Specifically, the abundance of 229 of the 313 identified mitochondrial proteins in soleus was significantly regulated in Anti-MuSK compared to Sham (FDR q < 0.05). The abundance of 172 out of 371 proteins was regulated in the diaphragm, and 11 out of 328 in the sternohyoideus muscle of Anti-MuSK rats. This pattern was consistent for proteins associated with the mitochondrial ribosome subpopulation (mtRibo, Figure 7C).

**Figure 7.**
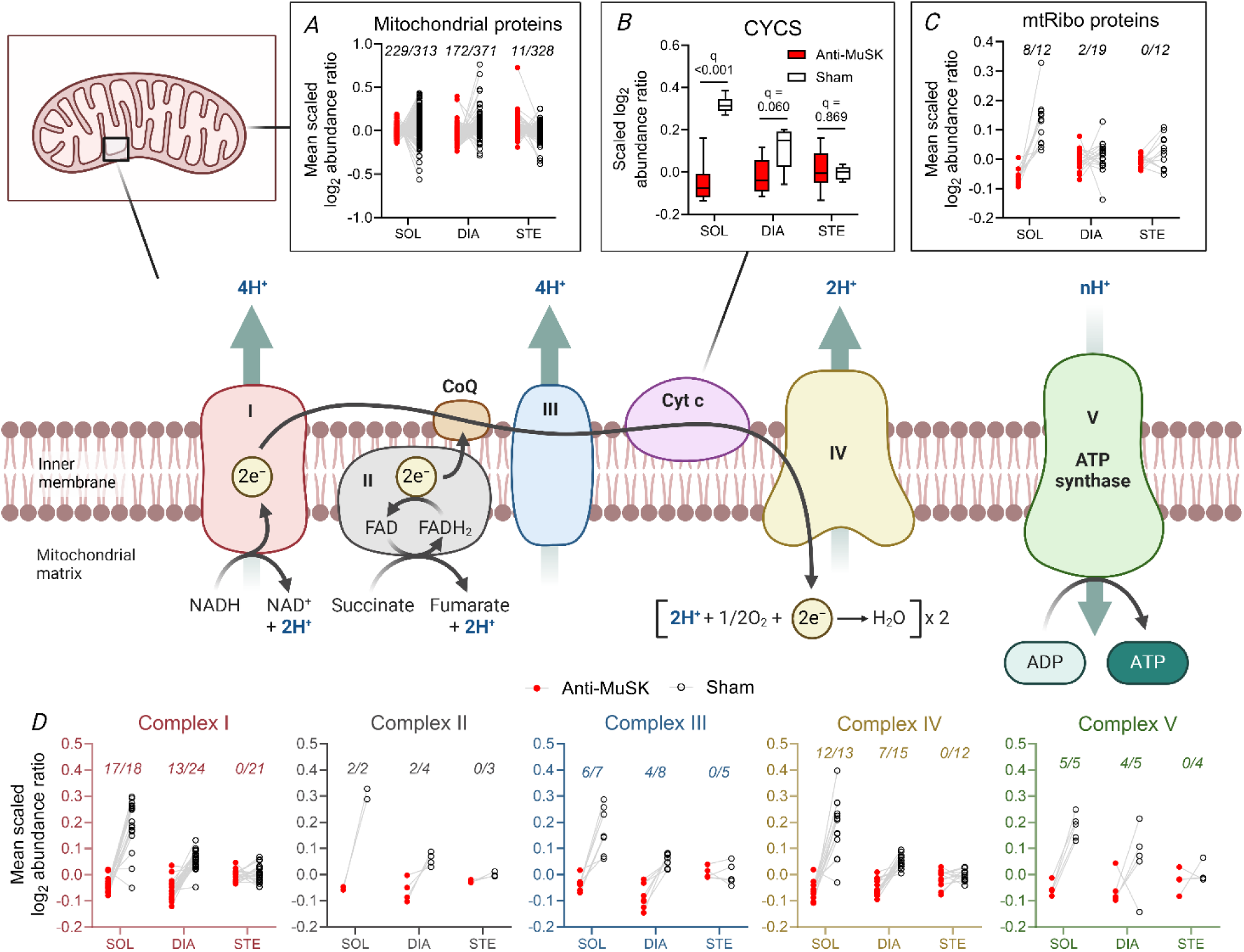
Regulation of mitochondria and oxidative phosphorylation (OXPHOS) in Anti-MuSK skeletal muscles. Panel A, C, and D depict the mean protein abundance of proteins within select mitochondrial categories extracted from the MitoCarta 3.0 catalog, with lines connecting the value between Anti-MusK (red circles) and Sham (open black circles). Panel A and C display data of all identified mitochondrial proteins within the MitoCarta 3.0 catalog and mitochondrial ribosomal proteins (mtRibo), respectively. Panel B displays the protein abundance of cytochrome C (CYCS), with values above box plots highlighting the FDR-corrected *P*-value (q-value) from the DEP analysis. Panel D displays abundance data from proteins in each electron transport chain/OXPHOS structure (complex I-V). For panels A, C, and D, the italic numbers above data points highlight the number of regulated proteins (FDR q <0.05), as revealed by DEP analysis, relative to the total number of identified proteins in the given muscle type (e.g., 17/18 indicates that a total of 17 out of 18 proteins are identified as either up- or downregulated). The figure was created based on a reprint of the “Electron transport chain” template retrieved from Biorender.com (2024).

Additionally, we observed a marked reduction in the abundance of protein components across complexes I-V of the electron transport chain (ETC), as well as of the nodal electron shuttler Cytochrome C in the soleus muscle (Figures 7B and 7D, FDR q < 0.05). A similar regulatory pattern was evident here, where the diaphragm muscle of Anti-MuSK samples was less affected than the soleus, while the ETC proteome of the sternohyoideus remained unaltered compared to Sham.

### Skeletal muscle myosin and ribosomal protein abundance

GSEA identified the myosin complex and ribosomal components as key regulated elements in Anti-MuSK skeletal muscles. For instance, the abundance of the slow-twitch/type I fiber-associated myosin heavy chain MYH7 was significantly downregulated in both the soleus and diaphragm muscles of Anti-MuSK-treated animals (Figure 8A, FDR q < 0.05). Similarly, several other myosin heavy chains (MYH6, MYH9, and MYH10) and myosin light chains (MYL2, MYL3, and MYL6) were downregulated in the Anti-MuSK soleus muscle compared to sham controls (FDR q < 0.05).

**Figure 8.**
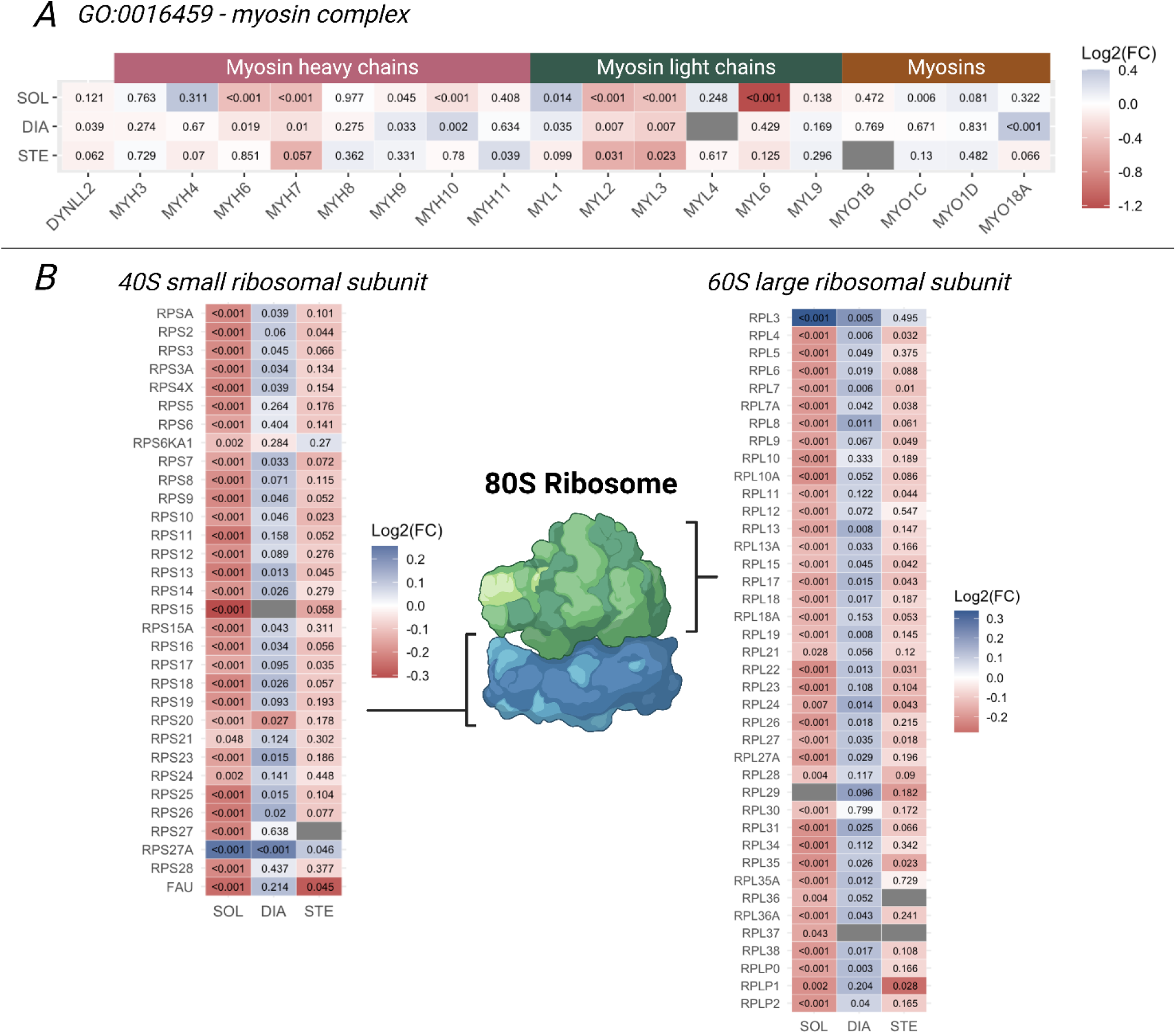
Myosin and ribosomal regulation in Anti-MuSK skeletal muscles. Panel A illustrates the identified proteins related to the myosin complex cellular component gene set (gene ontology GO:0016459), and panel B the identified cytosolic ribosomal proteins of the small 40S and large 60S ribosomal subunits that constitute the 80S eukaryotic ribosome complex. For both panels, the color of each heatmap tile indicates the log2FC (Anti-MuSK vs. Sham) per the scale bar to the right, and the number within each tile represents the FDR q-value derived from the DEP analysis. Grey tiles represent proteins not identified in the given TMT-plex run. For panel B, the figure was created based on a reprint of the “Eukaryotic ribosome” template retrieved from Biorender.com (2024). DIA, diaphragm muscle; SOL, soleus muscle; STE, sternohyoideus muscle.

In addition, the abundance of nearly all identified ribosomal proteins from the small 40S subunit was downregulated in the Anti-MuSK soleus muscle, except for RPS27A (FDR q < 0.05, Figure 8B). A similar pattern was observed for the large 60S ribosomal subunit, where all proteins except RPL3 were downregulated in the Anti-MuSK soleus muscle (FDR q < 0.05). Although to a lesser extent than in the soleus, the sternohyoideus muscle of Anti-MuSK animals also showed a reduction in the abundance of selected proteins from both the small and large ribosomal subunits (4 out of 31 and 11 out of 38, respectively). In contrast, the diaphragm muscle of Anti-MuSK animals exhibited an increase in the abundance of several ribosomal proteins from both the small and large ribosomal subunits (16 out of 31 and 25 out of 39, respectively).

## 4. Discussion

Since MuSK-MG was first identified as a MG subtype in the early 2000s, there has been a growing interest in a deeper myocellular characterization and basic disease understanding of the MuSK-MG phenotype. To this end, we aimed to characterize the skeletal muscle phenotype of MuSK-MG with special emphasis on the effect of the disease on skeletal muscle mass and its proteome signature. The main findings included distinct muscle type- and fiber type-dependent effects of the MuSK antibody attack on muscle mass as well as muscle-specific proteome alterations.

Providing proof of the active immunization procedure, MuSK antibody seropositivity was confirmed in all Anti-MuSK rats, and Anti-MuSK serum further demonstrated specific binding of rat IgG at the NMJ in wild-type soleus muscle. A clear neuromuscular histopathological finding was likewise revealed in Anti-MuSK rats, with the occurrence of fragmented and denervated NMJs compared to the muscle of sham controls. Moreover, soleus muscles exhibited marked atrophy in Anti-MuSK rats in concordance with the loss of body mass, while the mass of the EDL and gastrocnemius muscles remained unchanged compared to sham rats. Collectively, these findings confirmed the presence of both NMJ disruption and muscle atrophy in Anti-MuSK rats, validating the success of the active immunization procedure in inducing a preclinical disease phenotype resembling MuSK-MG, as previously reported by others (27).

To build on previous reports of muscle atrophy being a trait of MuSK-MG (5,8,17–21,23–28), we performed detailed analysis of the muscle atrophic phenotype in the Anti-MuSK rats. Among the different skeletal muscle types included, the slow-twitch fiber dominant soleus exhibited clear signs of atrophy in Anti-MuSK rats, while fast twitch dominant muscles, EDL and gastrocnemius, were unaffected. Moreover, soleus atrophy occurred proportionally to the loss of body weight in the Anti-MuSK rats, as indicated by a lack of difference in soleus muscle wet weights between groups when normalized to body weight, while the relative wet weight of both EDL and gastrocnemius was both proportionally higher in Anti-MuSK. It has been shown that marked extra-synaptic expression of MuSK is present in the soleus, but not TA muscle, in mice (26). This immediately suggests signal-transducing properties of MuSK beyond the NMJ, specifically in slow-twitch muscle. Further, recent reports indicate that MuSK functions as a co-receptor for promoting BMP signaling to engage specifically in mass regulation in slow-twitch muscle fibers (29,31). Specifically, inhibition of BMP-Smad signaling is believed to reduce the muscle hypertrophic response in myostatin-deficient mice, potentially by releasing the suppressive regulation upheld by ubiquitin ligase FBXO30 under normal conditions (58). Thus, antibody-mediated disruption of MuSK function triggers degradation signaling in slow-twitch fibers with concurrent loss of NMJ integrity across all fiber types. In our analysis of the fiber size of slow-twitch type I fibers in the TA, EDL, and sternohyoideus muscles, we observed atrophy exclusively in this fiber type.

To our knowledge, this is the first study to characterize the skeletal muscle proteome related to MuSK-MG. The three skeletal muscles selected for proteomics analysis represent distinct anatomical locations and functional roles in rodents. The soleus muscle represents a slow-twitch fiber dominant muscle involved in locomotion; the diaphragm, a primary respiratory muscle characterized by a more mixed distribution of slow-twitch and fast-twitch fibers, and; the sternohyoideus, an infrahyoid neck muscle involved in both respiration and swallowing characterized by a dominance of fast-twitch fibers (59,60). Differential protein expression analysis revealed varying degrees of regulation across these muscle types, with the greatest changes observed in the soleus and the least in the sternohyoideus. The extent of differential protein regulation between muscle types may reflect susceptibility of a specific muscle type to disuse-mediated atrophy due to the loss of NMJ integrity (61) as well as the differences in fiber type composition. We did not include the diaphragm in our analysis. However, we observed fiber atrophy in the relatively small slow-twitch fiber population of the sternohyoideus, despite a general lack of atrophy when judged on a whole-muscle level. This suggests that the proteomic changes identified in this study may result from a combination of disuse-mediated atrophy due to NMJ disruption and the effects of loss of MuSK and downstream signaling via e.g. BMP signaling (30).

Despite differences in proteome phenotype between the three analyzed muscle types, two proteins, NCAM1 and MUSTN1, were identified and ranked among the top upregulated proteins in all three muscle types in Anti-MuSK muscles compared to sham. Hence, their upregulation can be interpreted as a generic trait of the disease phenotype irrespective of the observed differential muscle atrophy. The neural cell adhesion protein NCAM1/CD56 has been proposed as a biomarker within neuromuscular diseases (e.g., Charcot-Marie-Tooth disease) and sarcopenia as increases in its expression may reflect NMJ instability and denervation (62,63). A previous study explored muscle NCAM1 expression in mice following passive immunization with MuSK IgG derived from human patient serum but reported no effects of the procedure in a small sample size (64). Recently, MUSTN1 has received attention as a protein secreted from smooth muscle and is suggested to be important for skeletal muscle regeneration in response to e.g., injury (65).

Analysis of the linear relationship between scaled protein abundance and soleus wet weight revealed multiple proteins with strong positive or negative correlations. For instance, among proteins of which the abundance strongly correlated negatively with soleus muscle mass (*r* < -0.9), proteins involved in ubiquitin- and ubiquitin-like-mediated protein degradation, namely UBE2N, NEDD4, and SUMO3, were identified. The well-described ubiquitin E3 ligase MuRF1, strongly associated with muscle atrophy, has recently been shown to engage in ubiquitylation with the ubiquitin-conjugating enzyme UBE2N (66). Our proteomics workflow also identified MuRF1/TRIM63 with high confidence in soleus and sternohyoideus, and for both muscle types it was upregulated in the Anti-MuSK phenotype (FDR q <0.05). However, the abundance of MuRF1 only displayed a vague negative correlation with soleus muscle mass (*r* = -0.16). Previous work has shown that passive immunization with patient-derived MuSK IgG may promote atrophy signaling in mouse skeletal muscle, as evidenced by increased MuRF-1 protein expression (64). Likewise, the HECT-domain ubiquitin ligase NEDD4 has also been demonstrated to be central in denervation-induced muscle atrophy (67). Finally, among proteins with strong negative associations between abundance and soleus muscle mass, many were related to calcium handling and signaling (e.g., CAMK2G, STIM1, CASQ1, CAPN1), glutathione metabolism, and histones. This pattern may reflect an increased demand for calcium handling due to compromised calcium storage and buffering, particularly within mitochondrial pools, as well as elevated calpain activity, which could constitute a proteolytic component to the observed muscle atrophy (68).

GSEA revealed an impact of the MuSK antibody attack on several myocellular components. For soleus, the mitochondrial and ribosomal proteomes as well as the myosin machinery were markedly impacted in the Anti-MuSK phenotype. At a smaller magnitude, a compromised mitochondrial proteome was also evident in the diaphragm of Anti-MuSK rats, whereas the type II fiber-dominant muscle, sternohyoideus, was unaffected. Accordingly, the mitochondrial deficits of several bioenergetic and structural properties (e.g., electron transport chain, mitochondrial ribosome, and inner and outer membrane) likely reflect a dysfunctional mitochondrial population in the MuSK-MG skeletal muscle phenotype. This may hold multiple myocellular implications such as impaired bioenergetics and calcium handling (69). In addition, components of the muscle ribosomal translational machinery were likewise compromised in Anti-MuSK soleus, and to some extent in sternohyoideus, as evidenced by a downregulated abundance of proteins of the small and large eukaryotic ribosomal subunits. Notably, the opposite was observed for ribosomal components in the diaphragm of Anti-MuSK rats, indicating an upregulated translational capacity in this muscle phenotype. This was accompanied by an increase in the abundance of proteasomal protein components in the diaphragm, indicating regulatory countermeasures to maintain muscle proteostasis in the diaphragm of the Anti-MuSK phenotype.

Despite our comprehensive methodological approach to characterize muscle mass and proteome of MuSK-MG, our study includes some potential limitations. We acknowledge that the scope of our muscle molecular findings is constrained by the cross-sectional and descriptive nature of this study. The rat model employed, using active immunization with a MuSK antigen, may also be viewed as somewhat rapid but transient, given the disease’s onset and progression. In rats that are not euthanized due to humane endpoint or scheduled termination, the disease phenotype of the model typically remains stable for approximately 3-4 weeks after symptom onset, followed by spontaneous remission (data not shown). Regarding our proteomics workflow, we identified approximately 2,000 proteins in each of the three muscle types analyzed, with findings and interpretations limited to the extent of this proteomic coverage. The wide dynamic range of the skeletal muscle proteome presents challenges for comprehensive coverage in muscle proteomics (70). However, advances in sample preparation techniques, LC-MS/MS technology, and computational proteomics workflows continue to enhance the depth of proteome coverage. Additionally, incorporating methods such as laser-capture microdissection or single muscle fiber preparations could improve spatial resolution, helping to e.g., pinpoint changes adhering to the NMJ or elucidate molecular mechanisms underlying fiber type-specific muscle wasting. Furthermore, the translational relevance of the molecular landscape observed in preclinical rodent MuSK-MG models to human patients requires further validation. Key findings such as fiber type-specific muscle atrophy, protein abundance patterns linked or unrelated to muscle atrophy, and the disrupted mitochondrial proteome in affected muscles should be confirmed in human patients. Such validation could help inform the development of future therapeutic strategies for MuSK-MG patients.

In summary, our findings underscore the significant negative influence of MuSK antibodies on molecular properties of importance for skeletal muscle homeostasis in this MuSK-MG disease model. Disruption of MuSK function appears to induce muscle atrophy in a fiber type-dependent manner, with associated proteome remodeling indicative of severe myopathy-like pathophysiology that coincides with the loss of NMJ integrity. The identification of several regulated myocellular components and individual proteins, both linked to and independent of muscle wasting in the MuSK-MG phenotype, calls for further investigation.

## Supplemental material & data availability

The raw mass spectrometry proteomics data is available at the ProteomeXchange repository PXD069671, PXD070377, and PXDXXXXXX.

## Author contributions

J.E.J., T.H.P., K.V., A.R., M.B.S., M.B.L, and N.H. conceived and designed the study. J.E.J., T.H.K., and P.J., performed sample preparations for proteomics. J.P. performed mass-spectrometry and proteomics database searches. J.E.J. performed proteomics data analysis and bioinformatics. P.B.T. performed immunohistochemistry staining, confocal microscopy and analyses related to NMJ morphology and serum staining. Related to muscle fiber size analysis, J.E.J. performed immunohistochemistry staining, J.W., performed immunofluorescence microscopy, and J.E.J., performed analyses. J.E.J wrote original manuscript draft with major revisions from K.V., J.P., and T.H.P. All authors provided review, input, and final approval of the manuscript.

## Funding support

The study was funded by the Innovation Fund Denmark (grant no. 2041-00011B, recipients J.E.J., K.V., M.B.S., A.R., and T.H.P.,), A.P. Møller Foundation (Fonden til Lægevidenskabens Fremme, grant no. L-2022-00234, recipient J.E.J.), Læge Sofus Carl Emil Friis og Hustru Olga Doris Friis’ Legat (grant no. FID4300092, recipient J.E.J.), and Helga and Peter Korning’s Fund (grant no. DC472123-007-42, recipient J.E.J.).

## Acknowledgments

Laboratory technician Randi Scheel (Dept. of Public Health, Aarhus University) is acknowledged for assisting with sample cryo-sectioning.

## Conflict of interest

The authors have declared that no conflict of interest exists.

## Notes

https://doi.org/10.6084/m9.figshare.31698931

https://doi.org/10.6084/m9.figshare.31698877

### Competing Interest Statement

The authors have declared no competing interest.

