## Supplemental figures S1-S3 for "The muscle atrophic phenotype of MuSK myasthenia gravis: Insights from a preclinical rat model"

**Supplementary figures**

S1 Immunohistochemical staining for serum MuSK antibody reactivity and neuromuscular junction (NMJ) morphology

S2 Networks of regulated biological processes (GOBP) in Anti-MuSK skeletal muscle

S3 Gene set enrichment analysis of Reactome and WikiPathways terms

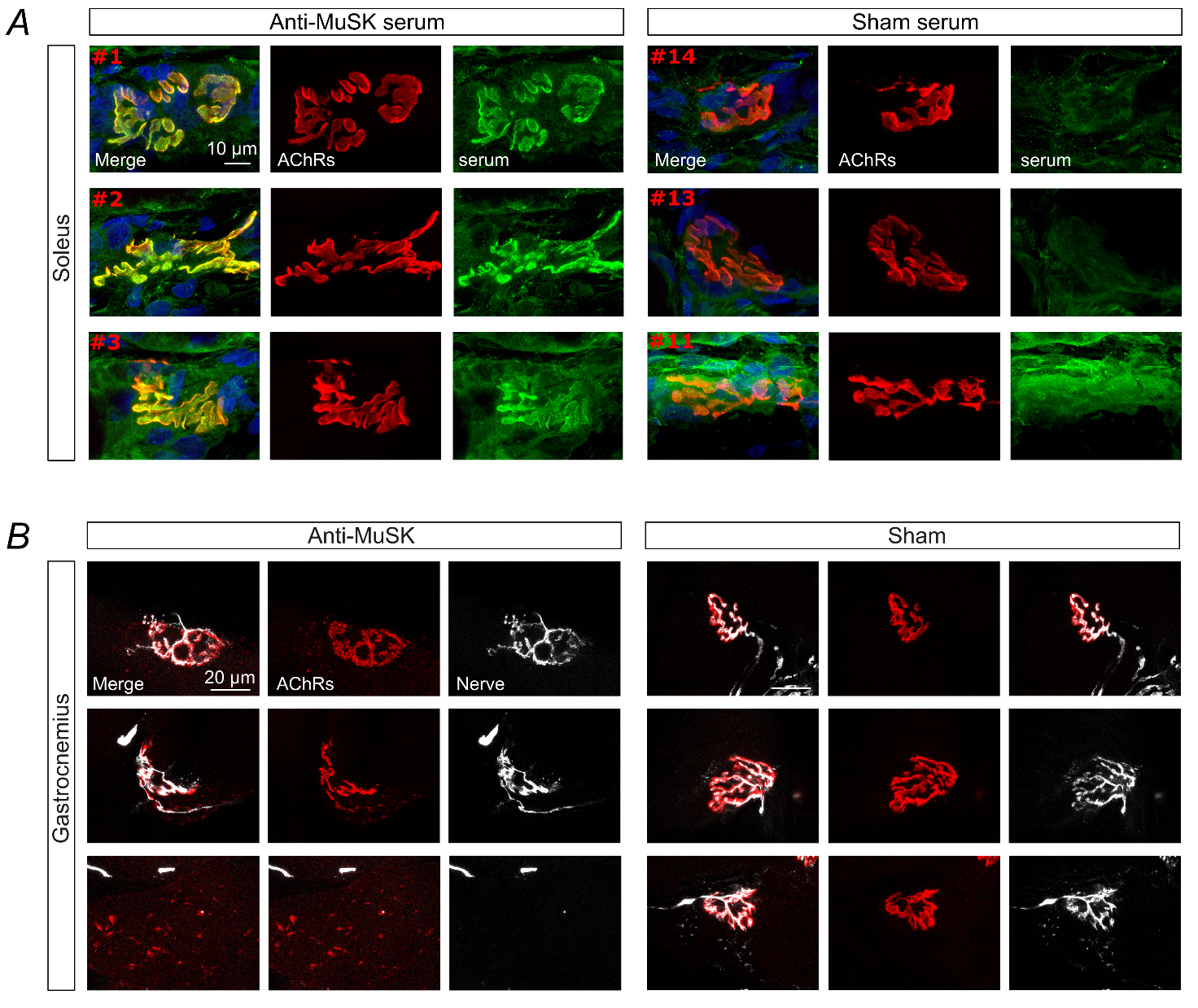

Figure S1 - Immunohistochemical staining for serum MuSK antibody reactivity and neuromuscular junction (NMJ) morphology. Panel A depicts representative confocal microscopy images confirming the localization of MuSK antibodies at the post-synaptic area in longitudinal cryosections from wild-type soleus muscles when incubated with serum collected from Anti-MuSK rats (left panel half). The right half of panel A illustrates only the background signal following incubation with Sham serum thus confirming the seronegativity of MuSK antibodies in the Sham rats. Red, bold numbers in the upper left corner of each merged image represent the animal ID. Anti-MuSK animals #1, #2, and #3 were selected based on Ab titer values representing a high-to-low range of titer (106.1, 80.7, and 22.7 nM, respectively). Panel B shows representative images of identified NMJs by colocalization of post-synaptic AChRs (α-bungarotoxin staining) and pre-synaptic nerve terminal markers SV2 and NF-M in gastrocnemius muscle from Anti-MuSK (left panel half) and sham (right panel half) rats. Histological findings of fragmented AChR clusters and decreased colocalization of pre- and post-synaptic markers (i.e., denervation) confirm the impact of MuSK antibodies induced by the active immunization with MuSK antigen thus mimicking the MuSK myasthenia gravis phenotype.

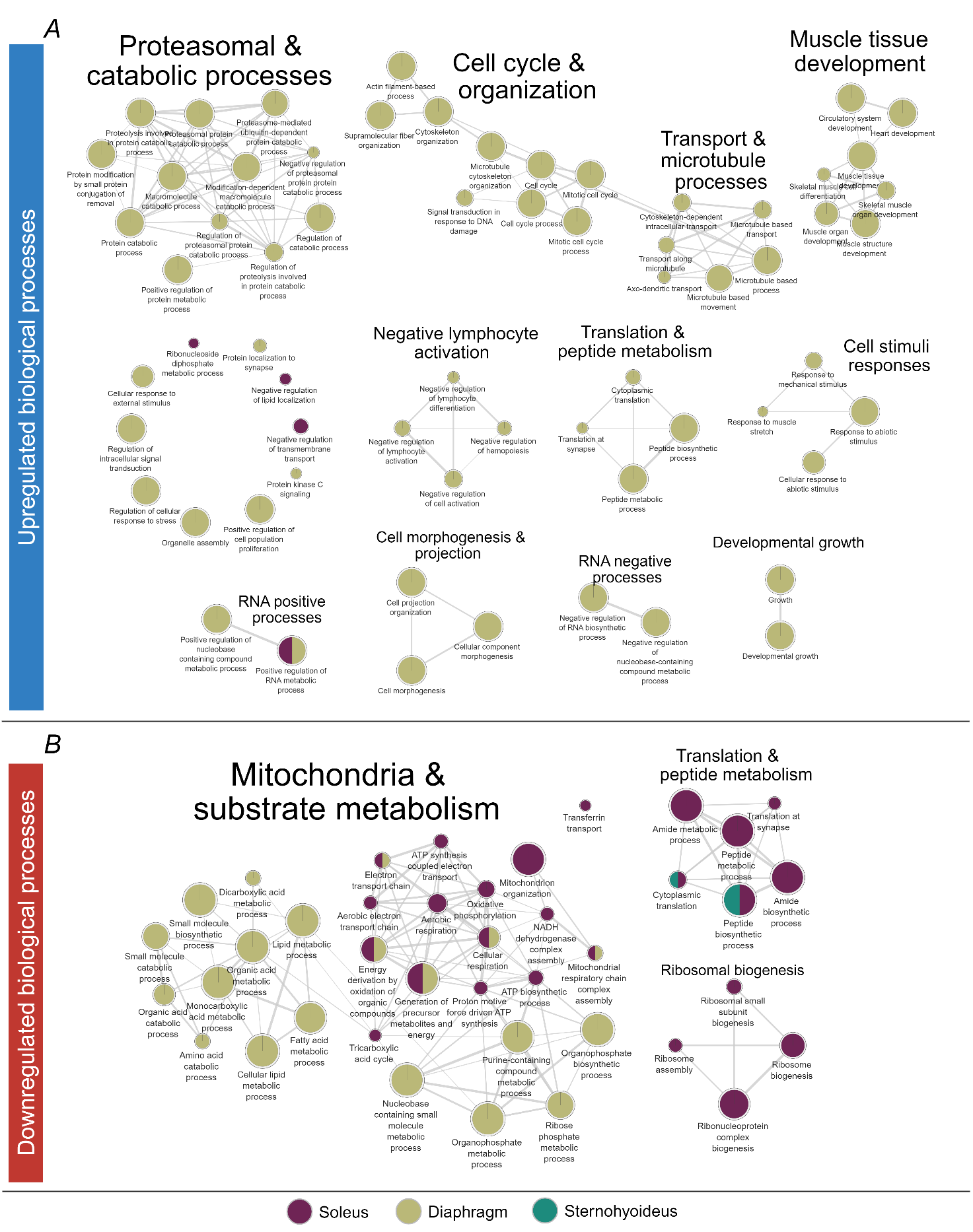

Figure S2 – Networks of regulated biological processes in Anti-MuSK skeletal muscles. Network visualization of up- and downregulated gene ontology biological processes (GOBP) are displayed in panels A and B, respectively. Networks were created using *Cytoscape* [S19] and the plugins of *EnrichmentMap* [S20] and *AutoAnnotate* [S21] by importing the results of the gene set enrichment analyses from pre-ranked proteins lists (based on π-value) on curated GOCC gene sets. Only enriched terms (FDR q <0.05) were used for visualization. The headline of each node network represents a broader annotation of the gene terms in the specific group. Each node represents a specific GOBP term with node size being the number of proteins in that term. Edge thickness between nodes represents the fraction of shared proteins between terms. Single nodes without edges represent significantly enriched terms with no shared proteins with other terms.

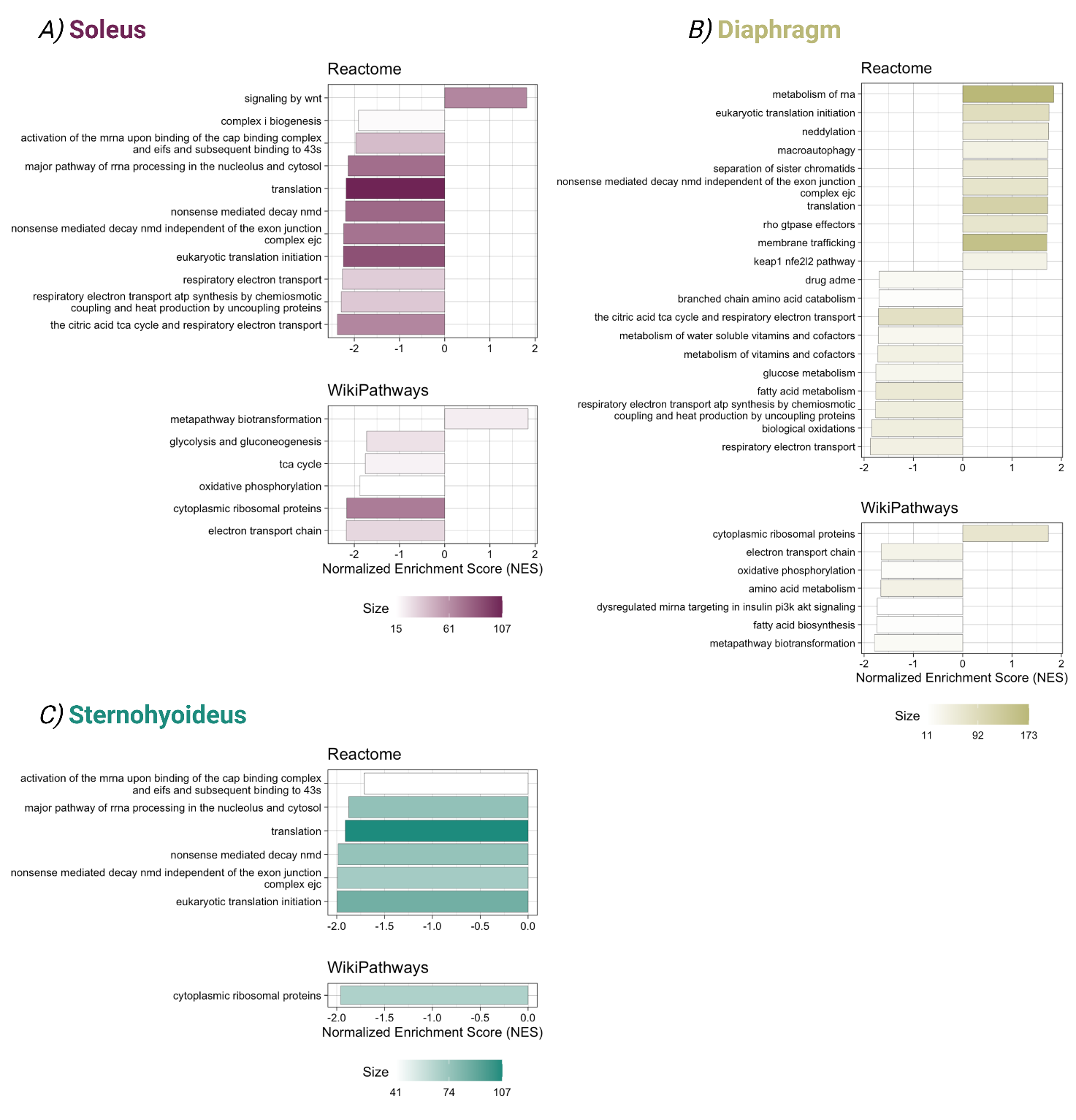

Figure S3 – Gene set enrichment analysis revealing regulated Reactome and WikiPathways terms in Anti-MuSK muscles. Results from gene set enrichment analysis based on pre-ranked protein lists (ranked by π-value) in soleus (A), diaphragm (B), and sternohyoideus (C) muscle using curated gene sets from the Reactome [S17] and WikiPathways databases visualized by horizontal bar graphs with only significantly enriched terms (FDR q <0.05). The x-axis indicates the direction of regulation as interpreted from the normalized enrichment score (NES), with >0 and <0 being terms significantly up- and downregulated, respectively. Bar colors depict the term size (number of proteins) as indicated by the scale bar.
